# Structure of a dimeric ABC transporter

**DOI:** 10.1101/2023.04.21.537812

**Authors:** Sarah C. Bickers, Samir Benlekbir, John L. Rubinstein, Voula Kanelis

## Abstract

ATP binding cassette (ABC) proteins generally couple ATP hydrolysis to the active transport of solutes across cellular membranes. All ABC proteins contain a core structure of two transmembrane domains (TMD1, TMD2) and two nucleotide binding domains (NBD1, NBD2), and many ABC proteins contain additional domains. Some members of the C subfamily of ABC (ABCC) proteins, such as the multidrug resistant protein 1 (MRP1), contain an N-terminal transmembrane domain (TMD0) and L0 linker that regulate transport activity and cellular trafficking, and mediate interactions with other proteins. Regulation can also be imparted by phosphorylation, proteolytic processing, and/or oligomerization of the proteins. Here we present the structure of yeast cadmium factor 1 (Ycf1p), a homologue of MRP1, in its mature form following cleavage by the yeast protease Pep4p. Remarkably, proteolytically cleaved Ycf1p forms a well-ordered dimer, with some monomeric particles also present in solution. Numerous other ABC proteins have been proposed to form dimers but no high-resolution structures have been reported. The monomeric and dimeric Ycf1p species are differentially phosphorylated at the intrinsically disordered regulatory (R) region, which links NBD1 to TMD2, and possess different ATPase activities indicating that dimerization affects the function of the protein. Protein-protein interactions involving TMD0, the L0 linker, and the R region mediate contacts between Ycf1p protomers in the dimer. In addition, cryo-EM density is observed for lipids at the interface between protomers, which suggests that lipids stabilize the dimer. The Ycf1p dimer structure is consistent with proposed dimerization interfaces of other ABCC dimers, such as MRP1.

## Introduction

ATP binding cassette (ABC) proteins are integral membrane proteins found in all kingdoms of life.^1-3^ ABC proteins contain a core structure comprised of two half-transporters, each containing a transmembrane domain (TMD) and a cytosolic nucleotide binding domain (NBD).^4-6^ Different ABC proteins are found either with the domains TMD1-NBD1-TMD2-NBD2 arranged in sequence in a single polypeptide (Fig. 1a), with separate polypeptides forming each half-transporter, or with separate polypeptides forming the TMDs and NBDs. Binding and hydrolysis of ATP at the NBDs of most ABC proteins powers the active transport of solutes across membranes, while in other ABC proteins ATP hydrolysis allows the protein to function as a channel or to regulate the activity of interacting proteins.^1,2^ Owing to these capabilities, ABC proteins are involved in numerous biological processes, such as nutrient import in prokaryotes, export of cytotoxic molecules in prokaryotes and eukaryotes, and peptide antigen presentation and cellular metal trafficking in eukaryotes.^1,2,7^

**Fig. 1.**
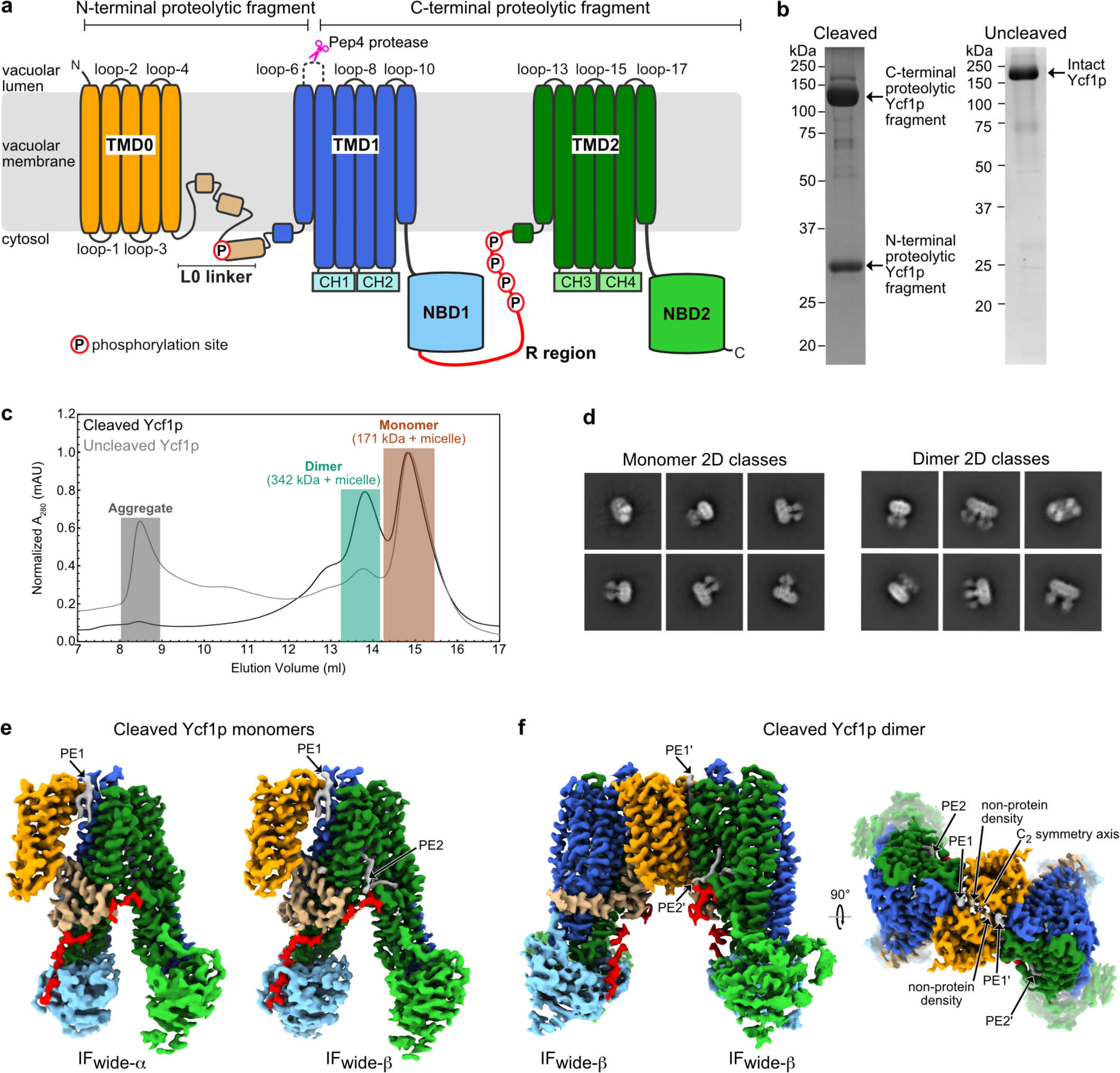
Structure of proteolytically cleaved Ycf1p. **a**, Schematic diagram of Ycf1p showing the three transmembrane domains (TMD0, TMD1, and TMD2), the L0 linker, the nucleotide binding domains (NBD1, NBD2), the regulatory (R) region, and the coupling helices (CH1, CH2, CH3, and CH4) that are located in TMD1 and TMD2 and contact the NBDs. Phosphorylation sites in the L0 linker (S251^34,98^) and R region (S903, S908, T911, and S914^33,35,36^) are depicted with a black “P” circled in red. The loop connecting the first two transmembrane helices in TMD1 (loop-6), which contains a 17-residue insertion compared to most other ABCC proteins, is shown by a dashed curve. Cleavage of this loop by Pep4p protease (pink scissors) yields the N-terminal and C-terminal proteolytic fragments.^61^ **b**, A 12% sodium dodecyl sulphate–polyacrylamide gel electrophoresis (SDS-PAGE) gel showing cleaved Ycf1p-3xFLAG. The two fragments of Ycf1p (measuring at ∼141 kDa and ∼33 kDa from the gel) are consistent with proteolytic cleavage of Ycf1p at the Pep4p site. **c,** A 12% SDS-PAGE gel showing a purified sample of uncleaved Ycf1p-3xFLAG. Migration of the band is consistent with the 173.8 kDa molecular weight of Ycf1p-3×FLAG. **c,** Analytical gel filtration chromatography of cleaved (black trace) and uncleaved (grey trace) Ycf1p. **d**, 2D class average images showing cleaved Ycf1p monomer (*left*) and dimer (*right*). **e**, Cryo-EM maps for two different conformations of the cleaved Ycf1p monomer in which the NBDs have different degrees of separation (IFwide-α and IFwide-β). The protein domains are colored as in panel **a** and the lipids are colored in grey. **f**, Cryo-EM map of cleaved dimeric Ycf1p, in which both protomers are in the IFwide-β conformation, with protein domains coloured as in panel **a** and lipids in grey.

Human ABC transporters are divided into seven subfamilies, designated as ABCA to ABCG, based on the primary sequence of the TMDs and NBDs in the core structure.^1,2^ While many ABC proteins contain only the core structure, some subfamilies of ABC proteins possess additional domains and linkers that regulate the activity of the protein.^3-5,8^ The ABCC subfamily contains the most functionally diverse set of proteins, and includes the multidrug resistant proteins (MRPs) that function as transporters, the cystic fibrosis transmembrane conductance regulator (CFTR) Cl^-^ channel, and the sulphonylurea receptors (SUR1, SUR2) that form the regulatory subunits in ATP sensitive K^+^ (K_ATP_) channels.^1^ All ABCC proteins contain an N-terminal extension composed of either TMD0 and an L0 linker (Fig. 1a) or just an L0 tail.^6^ ABCC N-terminal extensions are involved in cellular trafficking, mediate protein interactions, and/or regulate ABC protein activity.^9-15^ The TMD0 domain of ABC proteins is highly dynamic, and is usually absent or seen only at low resolution in cryo-EM structures of isolated ABCC (and ABCB) transporters.^13,15-22^ Many ABC proteins also possess disordered linkers, such as the one connecting NBD1 to TMD2 that is known as the regulatory (R) region in CFTR and other ABCC proteins (Fig. 1a).^4,5,23-26^

Regulation of the activity of ABC proteins can occur through several different processes. Phosphorylation, which generally occurs in loops, or in the R region or other intrinsically disordered regions, has been shown to alter the structural features and interactions of the disordered region and/or affect the activity of the protein. Regulation by phosphorylation has been observed for CFTR,^23-25,27-29^ SUR2A,^30,31^ MRP1,^32^ and Ycf1p,^33-36^ as well as ABC proteins from other subfamilies.^37^ ABC proteins have also been reported to oligomerize, including those from the ABCC subfamily.^10,38-58^ The function of oligomerization is not well understood but may provide an additional layer of regulation. Proteolytic processing is another regulatory mode and occurs in luminal loops in both human ABCA3^59,60^ and Ycf1p.^61,62^

Ycf1p is a yeast ABCC protein that transports glutathione-conjugated heavy metals such as Cd^2+^ from the cytosol into the vacuole to detoxify the cell.^63^ Pep4p protease-mediated cleavage of the luminal loop connecting the first two transmembrane helices in TMD1 (loop-6, Fig. 1a) yields mature Ycf1p and affects the substrate specificity of the protein.^61^ Here we present structures of the endogenous, Pep4p-cleaved Ycf1p. Surprisingly, proteolytic cleavage promotes the formation of a well-ordered dimer of the protein, which is held together by protein-protein interactions involving TMD0, the L0 linker, and the R region, as well as by protein-lipid interactions. The dimeric Ycf1p exists alongside the monomeric protein, with ATPase activity and phosphorylation decreased in the dimer compared to the monomer. The structures reveal the molecular features of dimeric ABCC transporters, and give insight into MRP1, a close homologue of Ycf1p that can functionally complement its activity.^64^

## Results and Discussion

### Proteolytically cleaved Ycf1p exists in multiple well-ordered oligomeric states

To obtain the endogenous proteolytically processed (or cleaved) Ycf1p, the W303-1A yeast strain, which expresses the Pep4p protease, was modified to include a 3×FLAG tag at the 3′ end of the *YCF1* gene in the chromosomal DNA. Similar to the procedure used to obtain uncleaved Ycf1p from a yeast strain lacking endogenous proteases,^35^ the modified yeast strain was cultured and harvested, and its membranes were isolated and solubilized with the detergent dodecyl-β-D-maltopyranoside (DDM). Ycf1p was then purified with anti-FLAG affinity chromatography and exchanged into the detergent glycol-diosgenin (GDN). Because the two proteolytic fragments of Ycf1p are metabolically stable and remain associated following cleavage by Pep4p,^62^ an intact but cleaved Ycf1p could be isolated by this procedure (Fig. 1b, *left* and Extended Data Fig. 1a, *left*). The resulting two fragments of Ycf1p have sizes consistent with proteolytic cleavage of loop-6, as described previously.^61^ The smaller fragment corresponds in size with Ycf1p TMD0, the L0 linker, and the first TM helix of TMD1 (calculated MW of 36.7 kDa *vs*. determined MW from SDS-PAGE of ∼33 kDa). The larger fragment is the size of the remaining TM helices 7-11 of TMD1, connecting loops, NBD1, R region, TMD2, and NBD2, as well as the 3xFLAG tag (calculated MW of 137.1 kDa *vs.* determined MW from SDS-PAGE of ∼141 kDa). Mass spectrometry of tryptic peptides confirms that the Pep4p cleavage site is located within loop-6, and specifically in the 17-residue insertion in loop-6 (Extended Data Fig. 1b).^61^ However, Pep4p-mediated cleavage of Ycf1p does not occur at a single site. Rather, the mass spectrometry data indicate multiple proteolytic sites between residues D325 and N335. In contrast, endogenous Ycf1p-3×FLAG from the modified BJ2168 yeast strain,^35^ which lacks the gene for Pep4p protease, is not proteolytically cleaved (Fig. 1b, *right* and Extended Data Fig. 1a, *right* and reference 35). Size exclusion chromatography of the proteolytically cleaved Ycf1p-3×FLAG, which will herein be referred to as cleaved Ycf1p, suggests multiple oligomeric states of the protein. However, while cleaved Ycf1p exists in both monomeric and dimeric forms (Fig. 1c, black trace), Ycf1p from a ΔPep4p strain appears as a primarily monomeric species along with some higher-order species (Fig. 1c, grey trace and reference 36). The fraction of aggregated protein cannot be discerned from these data, as aggregated protein scatters light and thus appears to absorb light strongly even at low concentrations.^65^

Cryo-EM of the cleaved Ycf1p (Extended Data Figs. 2, 3, and 4, and Supplementary Table 1) allowed calculation of two-dimensional (2D) class average images that show both monomeric Ycf1p (Fig. 1d, *left*), as well as dimeric Ycf1p (Fig. 1d, *right*). Image analysis yielded three-dimensional (3D) maps of monomeric Ycf1p in two different conformations (Fig. 1e and Extended Data Figs. 2, 3a-d, 4ab) and a 3D map of a well-ordered cleaved Ycf1p dimer (Figs. 1f and Extended Data Figs. 2, 3abe, 4c). The two maps of cleaved Ycf1p (at 3.4 Å and 3.1 Å) are in the inward-facing conformation with the substrate binding site exposed to the cytosol and the NBDs separated (Fig. 2 and Supplementary Table 1), as expected in the absence of bound nucleotide.^6^ These two different conformations of Ycf1p differ in the separation of their NBDs. Using nomenclature introduced previously, the inward-facing conformations of cleaved Ycf1p are referred to as IFwide-α and IFwide-β on account of their similarity to the IFwide, rather than the IFnarrow, conformation of uncleaved Ycf1p in the detergent digitonin.^36^

**Fig. 2.**
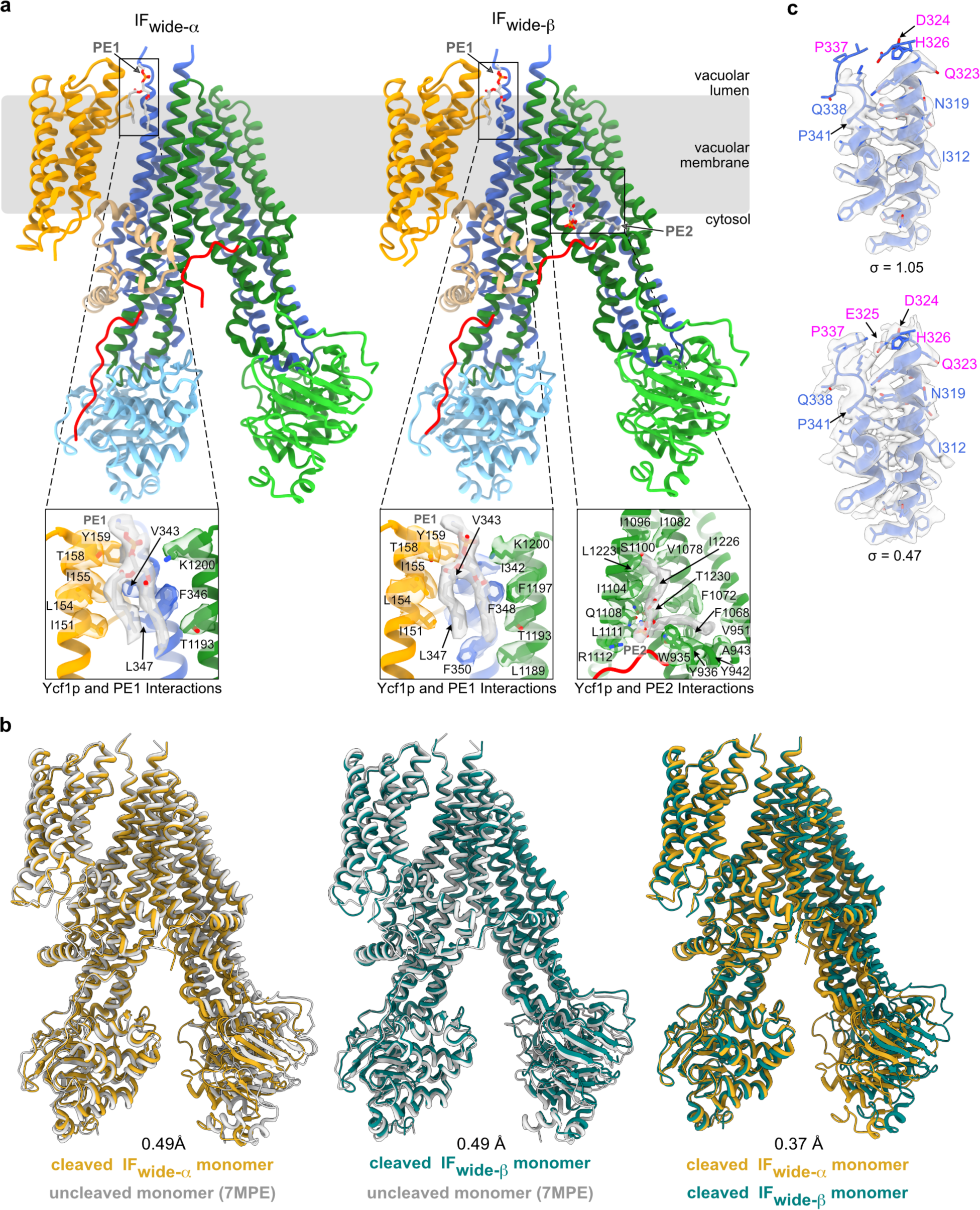
Lipid interactions in cleaved Ycf1p monomer and comparison with uncleaved Ycf1p. **a**, Atomic models of monomeric Ycf1p in IFwide-α (left) and IFwide-β (right) conformations, coloured as in Fig. 1. Close-up views of the interactions between lipids and each conformer are shown below. A phosphatidylethanolamine molecule (PE1) is located between TMD0 and TM helix bundle 1 on the luminal site of the protein for both the IFwide-α (*lower panel*) and IFwide-β (*lower left panel*) conformations and is shown as a stick in CPK colouring with the cryo-EM density in grey. The sidechains of interacting residues, defined as those within 4 Å of the lipid, are also shown as sticks with their cryo-EM densities. A phosphatidylethanolamine (PE2) is bound to the ABC core of the IFwide-β conformation (*lower right panel*). **b,** Overlay of the structure of uncleaved^35^ and cleaved Ycf1p monomers in the IFwide-α (*left*) and IFwide-β (*centre*) conformations. Overlay of the structures of the cleaved Ycf1p IFwide-α and IFwide-β monomers (*right*). The backbone of the uncleaved Ycf1p structure is in grey, while those of cleaved Ycf1p IFwide-α and IFwide-β monomers are in goldenrod and teal, respectively. **c,** Map-in-model fit for residues in TM6, TM7, and loop-6. The backbone is shown as a blue ribbon with all modeled sidechains shown as sticks. The cryo-EM density is shown at two different thresholds, denoted by the 0 values, to highlight the poor (or missing) density for loop-6. Residues labeled in pink are from the loop-6 insertion, whereas residues labeled in blue are from TM6, loop-6, and TM7, and are found in other ABCC proteins.

**Figure 3.**
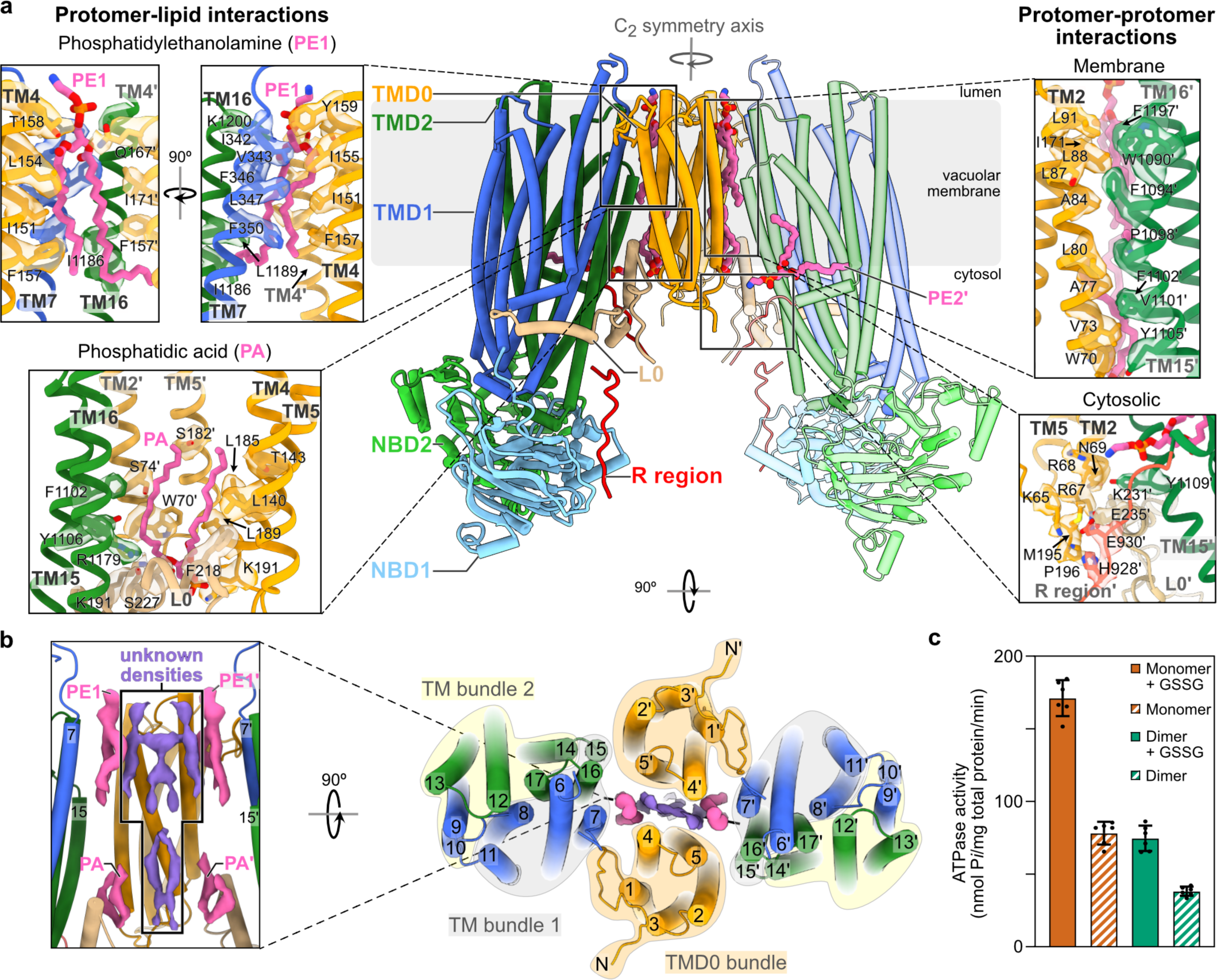
Structure and activity of cleaved, dimeric Ycf1p. **a**, An atomic model of the Ycf1p dimer is shown with cylinders representing α-helices and arrows representing β-strands (*centre*). The protein domains are coloured as in Fig. 1. One of the protomers (on the right-hand side) is coloured in lighter shades to distinguish it from the other protomer (on the left-hand side). Bound lipids that could be modeled are shown in CPK colouring except with pink instead of grey for the hydrocarbon tails. *Right*, Protein interactions between protomers are shown. The protein backbone is represented by a schematic ribbon diagram and interacting residues are shown as sticks along with their cryo-EM densities. *Left*, Protein-lipid interactions that link the two protomers are shown. Cryo-EM densities for the lipids are shown in Extended Data Fig. 7. **b,** The Ycf1p dimer is shown as viewed from the luminal side (*right*). *Left*, the density for additional hydrophobic molecules located at the TMD0/TMD0 interface between the two protomers is shown in purple. Cryo-EM densities for Ycf1p residues that interact with these molecules are shown in Extended Data Figure Fig. 7. **c,** Plots of the specific ATPase activity of monomeric and dimeric Ycf1p, without and with glutathione disulphide (GSSG).

CryoEM of dimeric Ycf1p shows the individual Ycf1p protomers in the dimer in the IFwide-α or the IFwide-β conformations, leading to three different dimer populations: IFwide-α:IFwide-α, IFwide-β:IFwide-β, and IFwide-α:IFwide-β (Extended Data Fig. 2). However, a high-resolution structure (at 3.2 Å) could be determined only for the IFwide-β:IFwide-β Ycf1p dimer (Fig. 1f and Extended Data Fig. 2). This map allowed construction of an atomic model for dimeric Ycf1p with both protomers in the IFwide-β conformation (Fig. 3 and Supplementary Table 1). As expected for ABCC proteins,^66^ TMD1 and TMD2 in the Ycf1p monomers and in each protomer in the Ycf1p dimer are domain-swapped such that TM bundle 1 and TM bundle 2 each possess TM helices from both halves of the ABC core (Extended Data Fig. 5).

As described above, size exclusion chromatography of purified uncleaved Ycf1p suggests the presence of some dimeric complexes in this sample also. In order to investigate if dimeric uncleaved Ycf1p was missed in the initial analysis of that sample,^35^ cryo-EM data for uncleaved Ycf1p was re-processed using the same workflow that led to a high-resolution structure of the cleaved Ycf1p dimer and showed some 2D classes consistent with a Ycf1p dimer (Extended data Fig. 6a). However, the protomers in this apparent dimer do not maintain consistent orientations relative to each other and 3D refinement of this dataset resulted in a low-resolution map with clear artifacts (Extended data Fig. 6b-d). Therefore, it appears that monomeric Ycf1p particles can interact with each other, at least transiently, before Ycf1p is cleaved, but a well-ordered and stable dimer structure forms only upon cleavage of the loop-6 insertion, allowing determination of the Ycf1p dimer structure.

### Nucleotide-free uncleaved and cleaved Ycf1p adopt similar conformations

Atomic models were constructed for both the IFwide-α and Ifwide-β conformations of monomeric cleaved Ycf1p, comprising ∼90% of the protein’s residues (Fig. 2a). Comparison of the cleaved IFwide-α and IFwide-β structures to three previously-determined structures of uncleaved Ycf1p^35,36^ indicates that cleavage of loop-6 (Fig. 1a) to yield the mature form of Ycf1p results in few structural changes in the protein (Fig. 2b). The r.m.s.d. values between the cleaved IFwide-α or IFwide-β conformer and the uncleaved Ycf1p determined in the same detergent system^35^ are both 0.49 Å when overlaying TM bundle 1 (Fig. 2b), which consists of transmembrane helices 6, 7, 8, 11, 15, and 16, and coupling helices (CH) 1 and 4 (Fig. 1a and Extended Fig. 5a). This lack of large-scale structural changes in the protein is consistent with the fact that loop-6 is disordered and does not interact with other regions of the protein, as evidenced by the lack density for most loop-6 residues in all inward-facing Ycf1p structures (Fig. 2c and references 35,36). Additionally, and as in the uncleaved Ycf1p structure,^35^ the structures of cleaved Ycf1p show a phosphatidylethanolamine molecule (PE1) nestled between TMD0 and TM helix bundle 1 on the luminal side of the protein, mediating additional contacts between TMD0, TMD1, and TMD2 (Fig. 1e and Fig. 2a, *lower panel* for IFwide-α and *lower left panel* for IFwide-β). The IFwide-β conformation also contains a phosphatidylethanolamine (PE2) contacting the C-terminal end of the R region and TMD2 (Fig. 1e, *right panel* and Fig. 2a, *lower right panel* for IFwide-β). Given that proteolytic processing alters the substrate specificity of Ycf1p,^61^ it is possible that there are differences in the uncleaved and cleaved Ycf1p structures at a different stage in the transport cycle, such as the substrate-bound state or when nucleotide is bound and the transporter is in the outward-facing conformation that allows the substrate to be released.

### Ycf1p dimerization involves TMD0, the L0 linker, and lipids

Examination of the Ycf1p dimer structure indicates that both protein-protein and protein-lipid interactions are critical for its formation (Fig. 3). The protein-protein interactions are formed by both membrane-embedded and cytosolic residues (Fig. 3a, *right panels*). Interactions within the membrane involve TMD0 residues from TM2 (*e.g.*, W70, V73, S74, A77, A84, L88, L91) and TM5 (I171′) from one protomer and TMD2 residues from TM15 (*e.g.*, W1090′, F1094′, P1098′, F1102′, Y1105′) and TM16 (F1197′) from the opposite protomer (residues in one protomer are labeled with “′” to distinguish them from residues in the opposite protomer). Note that in the SUR proteins, which possess an alternate orientation for TMD0 with respect to the ABC core compared to Ycf1p (and MRP1),^35^ TM2 of TMD0 binds TM helix bundle 1. Thus, the Ycf1p dimer structure, which has TM2/TM helix bundle 1 inter-protomer interactions in addition to TM4/TM helix bundle 1 intra-protomer contacts, further demonstrates the flexibility and plasticity of TMD0. Cytoplasmic loop interactions are formed between residues in the TMD0 loops (*e.g.,* K65, R67, N69) in one protomer and residues in the L0 linker (*e.g.,* K231′, E235′) and in the C-terminal end of the R region (*e.g.,* E928′, E930′) in the opposite protomer. Additional cytoplasmic contacts are made between the L0 linkers of each protomer (*e.g.,* M195, E235). While many of the cytoplasmic loop contacts involve polar and charged amino acids, both the polar groups of side chains (*e.g.,* the guanidino group of R67′ and carboxyl group of E235) and the non-polar groups of side chains (*e.g.,* aliphatic tails of R67′ and K231) mediate interactions.

Lipid molecules are also integral components of the dimer interface. As in both the monomeric cleaved Ycf1p structures presented here and a structure of uncleaved Ycf1p,^35^ a phosphatidylethanolamine molecule (PE1) is bound between TMD0 and TM bundle 1 on the luminal side of each protomer in the dimer (Fig. 3a, *upper left panels* and Extended Data Fig. 7). In addition to interacting with TMD0 and TM helix bundle 1 of one protomer, also observed in monomeric Ycf1p (Fig. 2 and reference 35), this phosphatidylethanolamine contacts TMD0 residues (F157′, Q167′, and I170′) in the opposite protomer in the Ycf1p dimer (Fig. 3a, *upper left panels* and Extended Data Fig. 7a, *right*). As observed for the cleaved Ycf1p IFwide-β monomer, each protomer of the Ycf1p dimer also contains a bound phosphatidylethanolamine (PE2) located where the C-terminal end of the R region meets TMD2 (Fig. 3a, *centre* and Extended Data Fig. 7). However, this lipid contacts the R region and TMD2 residues in one protomer only. Further, and as seen in monomeric Ycf1p structures, there are additional lipids bound to TMD1 and TMD2 (Extended Data Fig. 7, light purple density).

The Ycf1p dimer includes two lipid molecules, each of which is located on the cytoplasmic side of the protein and contacts both protomers (Fig. 3a, *lower left* and Extended Data Fig. 7b, *middle right*). These lipid molecules, which are modeled as phosphatidic acid (PA) owing to the lack of observable density for their head groups, contact TMD0 residues from both protomers (W70′, S74′, T143, S182′, L189). Each phosphatidic acid also binds the L0 linker (K191, F218, S219, S224, S227, K231) and TMD2 (F1102, Y1106, R1179) of one protomer. The cryo-EM map for the cleaved Ycf1p dimer also possesses non-protein density located on both the luminal and cytoplasmic sides of the dimer interface, between TMD0 of each protomer (dark purple density in Fig. 3b and in Extended Data Fig. 7). While the identity of these molecules is ambiguous, we hypothesize that they are also lipids based on cryo-EM density that is consistent with hydrocarbon lipid tails. These putative lipid molecules interact with multiple TMD0 residues in TM2 (W70, I71), TM4 (F147, L150, I151, L154, H157, T158), loop-4 (G161, W163, Q167), and TM5 (F170, I171, L174, F175, V177, I178, A181, S182, L186, L189), suggesting that they are important for the stability of the Ycf1p dimer (Extended Data Fig. 7b, *left panels*).

Studies of other α-helical transmembrane proteins indicate that the presence of lipids at protomer interfaces is inversely correlated with oligomeric stability,^67^ as assessed by buried surface area and number of electrostatic contacts between protomers. Further, for transmembrane proteins that depend on lipids for oligomeric stability, titration of the stabilizing lipid increases the stability and population of the transmembrane protein oligomer.^68^ Dimerization of Ycf1p results in only 1,613 Å^2^ of buried surface area and few electrostatic contacts, suggesting that the observed lipids are needed to stabilize the dimer. The limited buried surface area and electrostatic interactions also imply that the dimer interface is dynamic. Thus, it is possible that dimerization of Ycf1p will change with changes in lipid composition, which has been described for other transmembrane proteins.^67^ Further, changes in oligomerization may regulate the function of Ycf1p, as it does for other ABC transporters.^11,43,46,51-53,56,57^

### Dimeric cleaved Ycf1p is less active than monomeric cleaved Ycf1p

In keeping with a regulatory role for Ycf1p dimerization, the cleaved monomeric and dimeric Ycf1p species exhibit different levels of ATPase activity (Fig. 3c). Monomeric Ycf1p purified by gel filtration chromatography has a specific ATPase activity of 78.0 ± 6.0 nmol P*i*/mg total protein/min (mean ± s.d. from 3 independent assays from each of 2 separate protein purifications) compared to 38.0 ± 1.0 nmol P*i*/mg total protein/min (mean ± s.d. from 3 independent assays from each of 2 separate protein purifications) for dimeric Ycf1p purified by gel filtration. Considering that Ycf1p dimers are twice the molecular weight of Ycf1p monomers and that there are two Ycf1p molecules in each dimer, each protomer in the cleaved Ycf1p dimer has ∼2-fold lower ATPase activity compared to the cleaved Ycf1p monomer. This difference is found both in the presence and absence of the substrate glutathione disulfide (GSSG), which stimulates the ATPase activity of the monomer and dimer to the same extent. Based on this difference in ATPase activity, we checked if addition of MgATP or the non-hydrolyzable MgAMP-PNP alters the relative population of monomeric and dimeric Ycf1p species but found that it does not (Extended Data Fig. 8a), indicating that the Ycf1p dimer does not dissociate during the ATP hydrolysis reaction. One possible explanation for the difference in ATPase activities of the monomer and dimer is that the different species also possess different phosphorylation states. Mass spectrometry revealed that while S903, S908, T911, and S914 are all heavily phosphorylated in the cleaved Ycf1p monomer, only S903 is highly phosphorylated in the dimeric form of the protein (Extended Data Fig. 8b). Phosphorylation of S908 and T911 are required for Ycf1p transport activity,^33^ likely because of the increased ATPase activity of S908- and T911-phosphorylated Ycf1p.^36^ Thus, the higher ATPase activity of the Ycf1p monomer *vs.* the Ycf1p dimer is expected given the higher phosphorylation state of these residues in the monomer. Except for residues H928-K935, most of the R region is not observed in any of our cryo-EM maps of cleaved Ycf1p or is at low resolution and is modeled as poly-Ala. However, regardless of the level of phosphorylation, the R region in both the monomeric and dimeric cleaved Ycf1p structures binds the peripheral side of NBD1, which is opposite the nucleotide binding and NBD2 dimerization interface (red density in Fig. 1ef, and red curve in Figs. 2a and 3a). Interactions of the R region and the peripheral side of NBD1 are also observed in structures of uncleaved, phosphorylated Ycf1p (Fig. 4a*i* and references 35,36), albeit mediated by different R region residues in the different structures. The different R region interactions captured in the different uncleaved Ycf1p structures are likely because the intrinsically disordered nature of the R region allows for multiple NBD1/R region binding events.^35,36^ Previous work on Ycf1p^35^ and the ABCC protein CFTR^24,26,69-72^ show that the non-phosphorylated R region contacts NBD1 at the ATP binding and NBD2 dimerization sites (Fig. 4a*ii*), and that these interactions are disrupted with R region phosphorylation.^24,26,35,69,72,73^ At least for CFTR, this disruption leads to greater ATPase activity and ABC protein function.^27^ Thus, the non-phosphorylated S908, T911, and S914 and surrounding R region residues in the Ycf1p dimer, which is phosphorylated primarily at S903 only, may contact NBD1 in a manner that prevents ATP binding and NBD1/NBD2 dimerization, at least partly (Fig. 4a*iii*), leading to lower ATPase activity (Fig. 3c).

**Figure 4.**
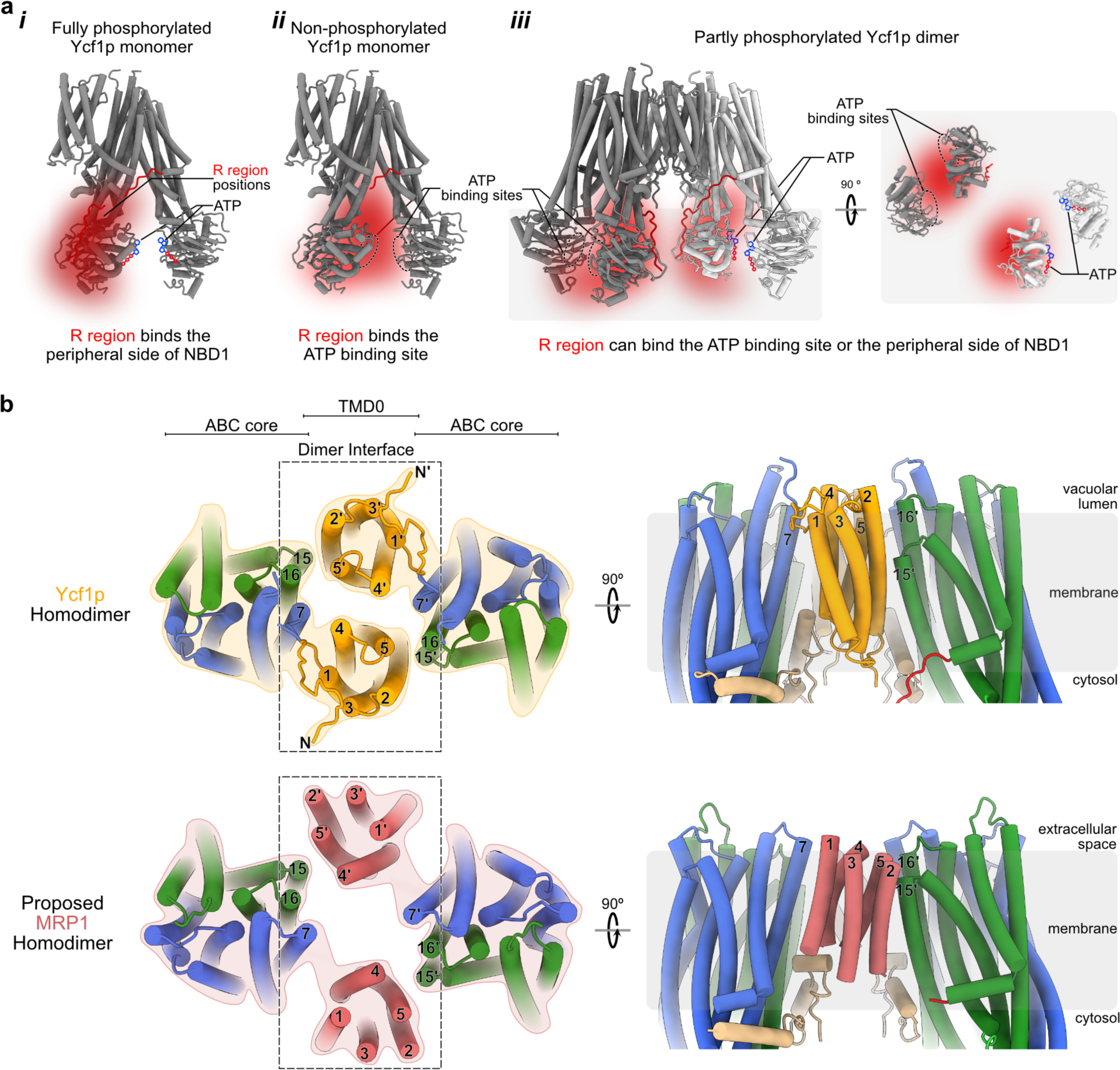
Models of R region interactions and other ABCC dimer structures. **a**, Schematic diagrams of the interactions of the R region with NBD1 in the fully phosphorylated Ycf1p monomer (*i*), the non-phosphorylated Ycf1p monomer (*ii*), and the partly phosphorylated Ycf1p dimer (*iii*). In each case, the possible locations for unobserved portions of the R region are shown by a diffuse red density to denote its multiple possible interactions, with the most likely locations of the R region in dark red and the less likely locations in light red. For clarity, the red diffuse R region density is labeled in panel *i* only. The fully phosphorylated R region likely binds the peripheral face of NBD1, allowing for ATP binding (*i*). ATP is shown with the adenine base and ribose in blue and the phosphates depicted as red circles. The non-phosphorylated R region likely interacts with the ATP binding site of NBD1, preventing nucleotide binding (*ii*), as depicted by the dashed ovals, which denote the ATP binding sites in NBD1 and NBD2 that overlay with the diffuse R region red density. Because structures of non-phosphorylated Ycf1p have not yet been determined, most of the R region is shown as a diffuse density. The partly phosphorylated R region in the Ycf1p dimer can interact with the ATP binding site or with the peripheral side of NBD1 (*iii*). ATP binds when the R regions binds the peripheral side of NBD1. **b,** Schematic diagram of the cleaved Ycf1p dimer (*top*) and the proposed MRP1 dimer (*bottom*). Two orientations are shown for each protein. The Ycf1p dimer is viewed from the lumen (*top left*) and from within the membrane (*top right*). Similarly, the MRP1 dimer is shown from the extracellular side of the plasma membrane (*bottom left*) and from within the membrane (*bottom* right). The TM domains of the cleaved Ycf1p dimer cryo-EM structure and the MRP1 dimer model are shown with cylinders representing α-helices. Transmembrane helices in TMD0 are coloured in orange for Ycf1p, as in Fig. 1a, and in coral for MRP1. As in Fig. 1a, transmembrane helices in TMD1 and TMD2 are in blue and green, respectively, for both proteins. Transmembrane helices in one protomer are labeled with “’” to distinguish them from those in the opposite protomer. While the structure of the ABC core is similar in Ycf1p and MRP1, the orientation of TMD0 helices differ slightly between the two proteins, leading to different inter-protomer TMD0 contacts between the Ycf1p and MRP1 dimers.

In addition to its transport function, Ycf1p recruits the soluble SNARE protein Vam7p to the vacuole during vacuolar fusion.^74^ This Ycf1p activity depends on a functional Walker A motif in NBD1, which binds ATP^6^ and which is occluded in the non-phosphorylated protein (Fig. 4a*ii* and reference 35). Notably, vacuolar fusion is also regulated by lipids, such as phosphatidic acid, and the lipid composition of vacuoles changes during the fusion process.^75-77^ Ycf1p has also been shown to interact with membrane and cytosolic proteins other than Vam7p.^78,79^ The oligomeric state of the protein, modulated by lipids or phosphorylation, could affect Ycf1p interactions and downstream functions.

### Ycf1p dimer structure is consistent with predicted dimers of other ABCC proteins

While high-resolution cryo-EM structures have been determined for MRP1 only in the monomeric state,^16-19^ studies of purified protein samples and studies in living cells indicate that MRP1 can also dimerize.^10,42,44,53^ The dimerization interface of MRP1 has been proposed to involve TMD0,^10^ and specifically that TM5 and the preceding extracellular loop (or loop-4 as defined in Fig. 1a) mediate the interactions between MRP1 protomers.^11^ Experiments also indicate that the L0 linker is involved in MRP1 dimerization,^11^ as with Ycf1p (Figure 3a, *lower right panel*). The Ycf1p dimer structure allows construction of a model of the MRP1 dimer that is compatible with these findings (Fig. 4b). This model of the MRP1 dimer was generated by aligning the ABC cores of two MRP1 proteins to the ABC cores of the two Ycf1p protomers in the Ycf1p dimer. Because the position of TMD0 relative to the ABC core differs slightly between these proteins,^35^ TM5 in MRP1 is at the dimer interface instead of TM2 in Ycf1p. Notably, and consistent with our finding of reduced ATPase activity for dimeric Ycf1p (Fig. 3c), co-expression of the N-terminal extension of MRP1 with full-length MRP1 reduces its transport activity.^10^

Similar to MRP1, only monomeric structures of CFTR have been determined previously.^70-73,80,81^ However there is substantial evidence that CFTR (or ABCC7) exists in both monomeric and dimeric forms in cells^41,43,45,47,48,53^ and that at least part of the CFTR dimerization interface is mediated by interactions between residues in the R region of each protomer.^47^ The fact that there are some R region/R region contacts between Ycf1p protomers in the dimer further establishes that the Ycf1p dimer structure is a good model for dimers of other ABCC proteins.

### Oligomerization interfaces vary between different classes of ABC transporters

ABC transporters outside the C subfamily have also been shown to oligomerize. Dimers and higher order oligomers have been detected in cells and in membrane preparations for the full-length transporters ABCA1,^46,51,56^ ABCA3,^57^ and P-glycoprotein (or ABCB1)^38-40^, as well as the dimeric half-transporter ABCG2.^49,50,52,55^ The oligomeric state of ABCA transporters can change during the transport cycle (*e.g.,* ABCA1)^51^ or based on the cellular location of the protein (*e.g.,* ABCA3^57^). Further, as seen for Ycf1p (Extended Data Fig. 8b), the phosphorylation state P-glycoprotein can differ in different oligomeric states.^40^ However, as with MRP1^16-19^ and CFTR^70-73,80,81^, high resolution structures have been determined only for monomeric states of ABCA1^82,83^, ABCA3^60^, P-glycoprotein,^84,85^ and the dimeric form of the half transporter ABCG2.^86^ The lack of dimeric structures for these ABC transporters, for which oligomerization has been detected biochemically, further suggests that oligomerization is transient or that the oligomeric state is unstable. In addition, ABCA1, ABCA3, ABCB1, and ABCG2 lack the N-terminal extension found in ABCC proteins and consequently ABCA, ABCB, and ABCG proteins must oligomerize differently than Ycf1p. Instead, ABCG2 has been suggested to oligomerize using TM helices and loops in TMD1,^52,55^ while ABCA1 dimerization employs the large extracellular domains^53^ that are hallmarks of the ABCA family.^4,6^

Multimeric complexes of ABC proteins are also found in prokaryotes, such as the complex between six copies of the ABC half-transporter HlyB, which are arranged as three HlyB homodimers, and three copies of the single pass transmembrane protein HlyD^58^. HlyB and HlyD form part of a type I secretion system in Gram negative bacteria.^87^ Similar to Ycf1p, associated lipids mediate critical interactions between the HlyB homodimers. However, and in contrast to Ycf1p, the HlyB transmembrane helices do not participate in complex formation and instead protein-protein contacts are formed between HlyB and HlyD. Further, and again unlike Ycf1p, the HlyB NBDs also form part of the interface between the HlyB homodimers. Because the Ycf1p dimer interface is partially formed by TMD0, the NBDs in different Ycf1p protomers are too far apart to contact one another in the inward-facing conformation seen here (Fig. 1, 3) and likely also in other states of the transporter.

## Conclusion

The studies presented here define the molecular structure of an ABCC transporter dimer. The dimeric protein displays different activity and phosphorylation compared to the monomeric form of the protein. Protein-protein contacts in the dimer involve the variably-oriented TMD0^35^ and the intrinsically disordered R region,^35,36^ highlighting the dynamic nature of ABC oligomerization. The structure also shows that bound interfacial lipids are likely integral to the stability of the Ycf1p dimer. These findings emphasize the flexibility of ABC dimers and explain why both monomeric and dimeric Ycf1p species are present in our preparations and likely also in cells, as seen for other ABC transporters.^38,39,41-43,45-49,51,53-56^ The Ycf1p dimer structure provides a model for understanding dimerization, and regulation by oligomerization, in other ABCC transporters.

## Experimental Procedures

### Construction and growth of the Ycf1p-3×FLAG Yeast Strain, and purification of Ycf1p-3×FLAG

The *S. cerevisiae* strain W303-1A (*MAT*a {*leu2-3,112 trp1-1 can1-100 ura3-1 ade2-1 his3-11,15*}) was further modified by insertion of sequence encoding a C-terminal 3×FLAG tag and a *URA3* marker downstream of the *YCF1* gene, as done previously in the BJ2168 strain ^35^. The modified W303-1A was name ySCB2.

Growth of the ySCB2 yeast strain and subsequent purification of cleaved Ycf1p-3×FLAG protein followed the procedure used for purifying uncleaved Ycf1p-3×FLAG from the strain, ySCB1 ^35^ with minor modifications. An 10 or 11-L culture of the ySCB2 yeast strain was grown in YPD medium (1% [w/v] yeast extract, 2% [w/v] peptone, 2% [w/v] dextrose) supplemented with 100 mg/mL ampicillin and 0.02% (v/v) antifoam, at 30 °C in a New Brunswick BioFlow fermenter. The culture was grown for a total of ∼48 h to an optical density at 660 nm (OD_600_) of 6.0 - 10.0. Yeast cells were pelleted by centrifugation at 4,000 × g for 15 min at 4 °C and resuspended in 1 mL of lysis buffer (8 g/L NaCl, 0.2 g/L KCl, 1.44 g/L Na_2_HPO_4_, 0.24 g/L KH_2_PO_4_, 80 g/L sucrose, 20 g/L D-sorbitol, 20 g/L D-glucose, 5 mM 6-aminocaproic acid, 5 mM benzamidine, 5 mM EDTA, and 10 mg/L PMSF at pH 7.4) per gram of cells. The resuspended yeast cells were lysed with a Biospec Beadbeater at 4 °C, with 0.5 mm glass beads in 5 cycles of 1 min on, 2 min off. Cellular debris was removed by centrifugation at 4,000 × g for 15 min at 4 °C, after which the membranes were collected via ultracentrifugation (Beckman L-90K) at 152,947 × g for 40 min at 4 °C. Membranes were resuspended, using either a Dounce or tissue homogenizer, in 0.5 mL of lysis buffer per gram of harvested cells and then frozen in liquid nitrogen and stored at −80 °C.

Frozen membranes were thawed in a tepid water bath and solubilized with 1% (w/v) n-dodecyl-β-D-maltopyranoside (DDM; Anatrace) with mixing for ∼1 h. Solubilized membranes were ultracentrifuged at 181,078 × g for 70 min (Beckman L-90K) and the supernatant containing Ycf1p was filtered through a 0.45 μm syringe filter (Pall) and applied to a 1.5 mL M2 affinity gel column (Millipore Sigma) that was pre-equilibrated with DTBS (50 mM Tris HCl, 150 mM NaCl, 0.04% [w/v] DDM, pH 8.0). The column was washed with 12 column volumes of DTBS, followed by 8 column buffers of GTBS (50 mM Tris HCl, 150 mM NaCl, 0.006% [w/v] glyco-diosgenin [GDN; Anatrace], pH 8.0). Ycf1p was eluted with 6 column volumes of GTBS containing 150 μg/mL 3×FLAG peptide. An additional five column volumes of GTBS without 3×FLAG peptide was applied to wash residual Ycf1p from the column. Elution fractions containing Ycf1p were concentrated with a 100-kDa molecular weight cut off (MWCO) Amicon Ultra centrifugal filter (Millipore Sigma). Samples for cryo-EM were concentrated to ∼5 mg/ml and used immediately, while samples for biochemical assay were concentrated to 1.2 - 2.2 mg/ml, flash-frozen in liquid nitrogen, and stored at −80 °C until use. When gel filtration chromatography was used, purified Ycf1p samples (100 μL) were loaded onto a Superose 6 Increase (300/60L) column that was pre-equilibrated with GTBS. The column was run at a flow rate of 0.1 mL/min at 4 °C using an ÄKTA Pure system. Where indicated, samples of purified cleaved Ycf1p were pre-incubated with 2 mM MgCl_2_ and either 2 mM ATP or AMP-PNP before applying the sample to the column.

### Malachite Green Assay

ATPase activity of Ycf1p was measured using the Malachite Green assay,^88^ which detects the amount of inorganic phosphate in solution. The cleaved Ycf1p monomer and dimer were obtained from size exclusion chromatography and used fresh in the ATPase assay. Samples of 20 µL that contained either 0.114 µg of the cleaved Ycf1p monomer or dimer in 50 mM HEPES (4-(2-hydroxyethyl)-1-piperazineethanesulfonic acid) pH 8.0, 3 mM MgCl_2_, 0.006% (w/v) GDN, and 2 mM ATP were prepared. The ATPase activity in the presence of substrate was measured by including 100 μM glutathione disulfide (GSSG), which has been used previously as a Ycf1p substrate.^36^ Control samples in the reaction buffer, with and without 100 μM GSSG, were prepared and phosphate standard samples (from 0 – 40 μM) were generated from a phosphate stock (Sigma). All samples were incubated at 30 °C for 90 min, then promptly diluted 8-fold in MilliQ H_2_O and flash frozen in liquid nitrogen until analysis.

Samples were thawed at room temperature for ∼20 min and 80 µL of each reaction were transferred to a 96-well plate. Each solution was then mixed with 20 µL of the phosphate detection reagent (0.01% [w/v] Malachite Green, 0.17% [v/v] Tween 20, 1.48% [w/v] ammonium molybdate) and incubated at room temperature for 20 min. The colour development reaction was quenched with 11 µL of a 34% (w/v) sodium citrate, the samples were mixed, incubated for 7 min at room temperature, and the absorbances measured at 620 nm with a SpectraMax i3X Multi-Mode Assay Microplate Reader. The amount of phosphate produced was determined based on a standard curved using the phosphate standards. Each measurement of phosphate production was corrected for the amount of phosphate produced in a sample that contained all components except the protein. ATPase assays were performed in triplicate for each of two independent cleaved Ycf1p preparations.

### Mass spectrometry analysis of Pep4p cleaved of Ycf1p

In-gel trypsin digestion of the two SDS-PAGE bands (at ∼32 kDa and ∼140 kDa) from purified Ycf1p samples followed by MS/MS was performed by SPARC Molecular Analysis Laboratory at The Hospital for Sick Children. Briefly, 10 μl of ∼0.35 mg/ml purified Ycf1p was loaded onto a 12% SDS-PAGE acrylamide gel. The gel was stained with Coomassie Brilliant Blue and destained with MilliQ. After dehydrating the gel bands with acetonitrile, the Ycf1p fragments were reduced by incubation with 10 mM dithiothreitol (DTT) at 60 °C for 1h and alkylated by incubation with 55 mM iodoacetate at room temperature for 20 min in the dark. The gel bands were re-dehydrated with acetonitrile, and the Ycf1p fragments were subsequently digested with trypsin (0.5 μg) at 37 °C overnight. The resulting peptides were subjected to LC MS/MS. The raw MS files were analyzed with Proteome Discoverer (v2.5.0.4.00) software using the *S. cerevisiae* database and the sequence of Ycf1p-3×FLAG protein. The parent and fragment mass tolerances were set to 10 ppm and 0.02 Da, respectively, and only tryptic peptides with a maximum of three missed cleavage sites were included. The addition of a carbamidomethyl group on Cys was specified as a fixed modification, while deamidation of Asn and Gln, oxidation of Met, acetylation of the N terminus, and phosphorylation of Ser, Thr, or Tyr were set as variable modifications.

### Mass spectrometry analysis of phosphorylation of Ycf1p

Similar to identification of the Pep4p cleavage site, in-gel trypsin digestion followed by MS/MS was used to analyze phosphorylation states in the cleaved Ycf1p monomer and dimer. However, in this case, purified Ycf1p was subjected to 4-16% Bis-Tris blue native PAGE. Gel bands corresponding to the cleaved Ycf1p monomer and the cleaved Ycf1p dimer were excised, dehydrated with acetonitrile, and incubated with DTT and iodoacetamide, as above. Following further dehydration of the gel bands, the in-gel cleaved Ycf1p monomer and dimer proteins were digested with trypsin and resulting peptides analyzed by LC MS/MS using the parameters described above. Phosphorylated and non-phosphorylated peptides were detected for both the cleaved Ycf1p monomer and dimer, and the ratio of the number of fragments detected from the phosphorylated and non-phosphorylated peptides, when the total number of fragments is high, approximately indicates the extent of phosphorylation.^35^

### Cryo-EM Sample Preparation and Data Collection

Freshly purified Ycf1p from the ySCB2 strain was applied to homemade nanofabricated holey gold grids^89^ with ∼1.2 µm holes. Grids were glow-discharged in air (Pelco easiGlow) for 2 min prior to the application of ∼1.5-3 µL of sample. Samples were frozen with a Leica Automatic Grid Plunging device with 1.5 s blotting time at 4 °C with a relative humidity of ∼100%. A total of 6,195 images were collected on a Titan Krios G3 electron microscope (Thermo Fischer Scientific) operating at 300 kV with a prototype Falcon 4i camera. Data collection was monitored using cryoSPARC Live^90^ and automated with the EPU software package. Two sets of movies were recorded. One set was recorded for 6.97 s, with a total exposure and exposure rate of 46.6 e/Å^2^ and 6.89 e/pixels/s, respectively. The second set of movies was recorded for 8.94 s, with a total exposure of and exposure rate of 46.0 e/Å^2^ and an exposure rate of 5.89 e/pixels/s.

### Imaging Analysis and Model Building/Validation

CryoSPARC v2^90^ was used post-data collection for image processing. Patch motion correction was applied to the first 29 exposure fractions of each movie and the contrast transfer function (CTF) of the resulting aligned images were estimated with patch CTF estimation. Movies were manually curated removing exposures with undesirable ice thickness, poor CTF fits, and sample aggregation, which resulted in a total of 6,195 movies used in particle analysis. Blob selection with particle dimensions between 250 and 350 Å was employed to generate initial 2D classes of monomeric and dimeric Ycf1p particles to use as templates for particle selection. Initial rounds of 3D particle sorting as well as non-uniform refinement^91^ were performed to remove non-Ycf1p particles and separate the monomeric and dimeric Ycf1p particles. Local motion correction^92^ was then applied to improve particle alignment and further rounds of 3D particle sorting and non-uniform refinement were conducted. 3D variability analysis^93^ was used to identify different modes of nucleotide binding domain (NBD) motion and generated templates with varying NBD distances, which were used in 3D sorting of conformations of both monomeric and dimeric Ycf1p. Final rounds of 3D particle sorting and non-uniform refinement provided five final cryo-EM maps (Extended Data Fig. 2). Cryo-EM maps for monomeric Ycf1p in the IFwide-α and IFwide-β conformations were at 3.4 Å and 3.1 Å, respectively. A cryo-EM map at 3.2 Å was obtained for a Ycf1p dimer in the IFwide-β:IFwide-β conformation. In contrast, maps of Ycf1p dimers in the Ycf1p IFwide-α:IFwide-α and IFwide-α:IFwide-β conformation are at 6.3 and 6.8 Å resolution, respectively. Models for the Ycf1p IFwide-α and IFwide-β monomers, and for the IFwide-βIFwide-β Ycf1p dimer were built in Coot^94^ using PDB 7MPE^35^ as a starting model. On account of the lower resolution of the NBDs and part of the R region, these regions were modeled as poly-Ala. Refinement was performed in both Phenix^95,96^ and iSOLDE^97^. Lipids were added using the Coot ligand library^94^ or imported using the SMILES string.

### Data deposition

EM maps for the cleaved Ycf1p monomer, in IFwide-α and IFwide-β conformations, and the cleaved Ycf1p dimer have been deposited in the Electron Microscopy Data Bank (EMDB), https://www.ebi.ac.uk/pdbe/emdb/ (EMDB accession codes: EMD-XXX, EMD-XXX, and EMD-XXX, respectively). The coordinates have been deposited in the Protein Data Bank, www.wwpdb.org (PDB ID code: YYYY, YYYY, and ZZZZ, respectively).

## Acknowledgements

SCB was supported by a Queen Elizabeth II Graduate Scholarships in Science and Technology and an Alexander Graham Bell Canada Graduate Scholarship-Doctoral from the Natural Sciences and Engineering Research Council of Canada (NSERC). JLR was supported by the Canada Research Chairs program. This research was supported by NSERC Discovery Grants to VK (RGPIN-2020-05835) and JLR (RGPIN-2023-04676). CryoEM data were collected at the Toronto High-Resolution High-Throughput cryoEM facility, supported by the Canada Foundation for Innovation and Ontario Research Fund.

## Statements of Contributions

VK conceived of the project and designed experiments with JLR and SCB. SCB created the yeast strain, purified the protein, performed ATPase assays, fabricated the cryo-EM specimen grids, processed cryo-EM images, and constructed the Ycf1p atomic models. SB prepared cryo-EM specimens and collected images. Figures were prepared by SCB and VK, and the manuscript was written by SCB, VK, and JLR.

