## Supplementary Figures for "Structure of a dimeric ABC transporter"

### Extended Data

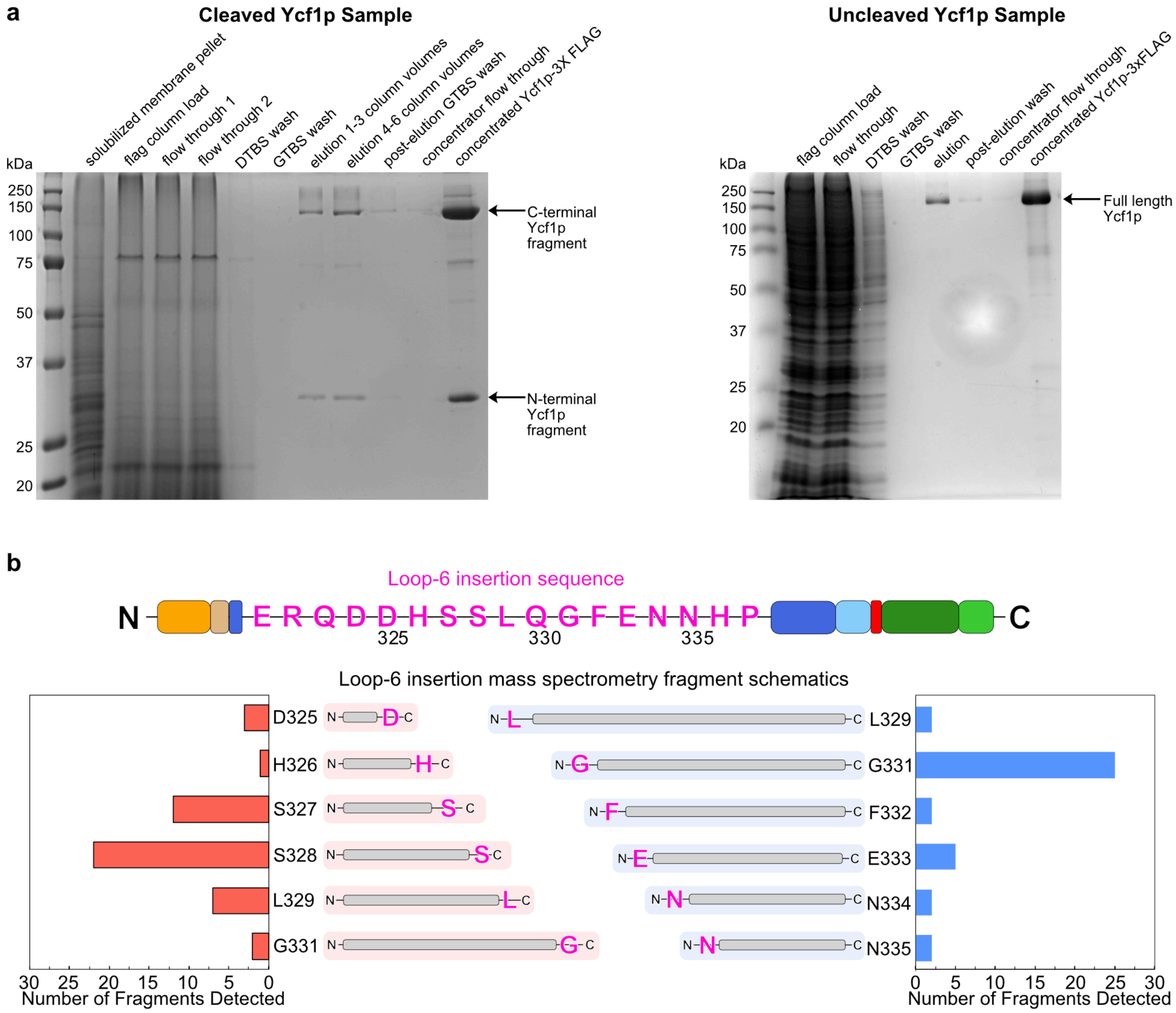


**Extended Data Fig. 1. Preparation and analysis of Ycf1p samples for cryo-EM. a**, 12 % SDS-PAGE of the affinity purification of cleaved Ycf1p-3xFLAG (*left*) and uncleaved Ycf1p-3xFLAG (*right*) for comparison. **b**, Proteolytic digestion occurs at multiple sites in the loop-6 insertion, the sequence of which is shown in pink (*top*). Histograms (*bottom*) displaying the number of fragments detected when the loop-6 residue is at the C terminus (*left*, red bars) or N terminus (*right*, blue bars). Each bar is labeled with a schematic of the corresponding fragment generated from MS/MS analysis of in-gel trypsin digestion of the N-terminal and C-terminal fragments of cleaved Ycf1p. Because SDS-PAGE bands for both the N-terminal and C-terminal fragment were subjected to trypsin digestion and subsequent MS/MS analysis, peptides that end either at loop-6 (red shading) or that begin at loop-6 (blue shading) are detected. The loop-6 residue at the C-terminal or N-terminal end of the peptide is in pink.

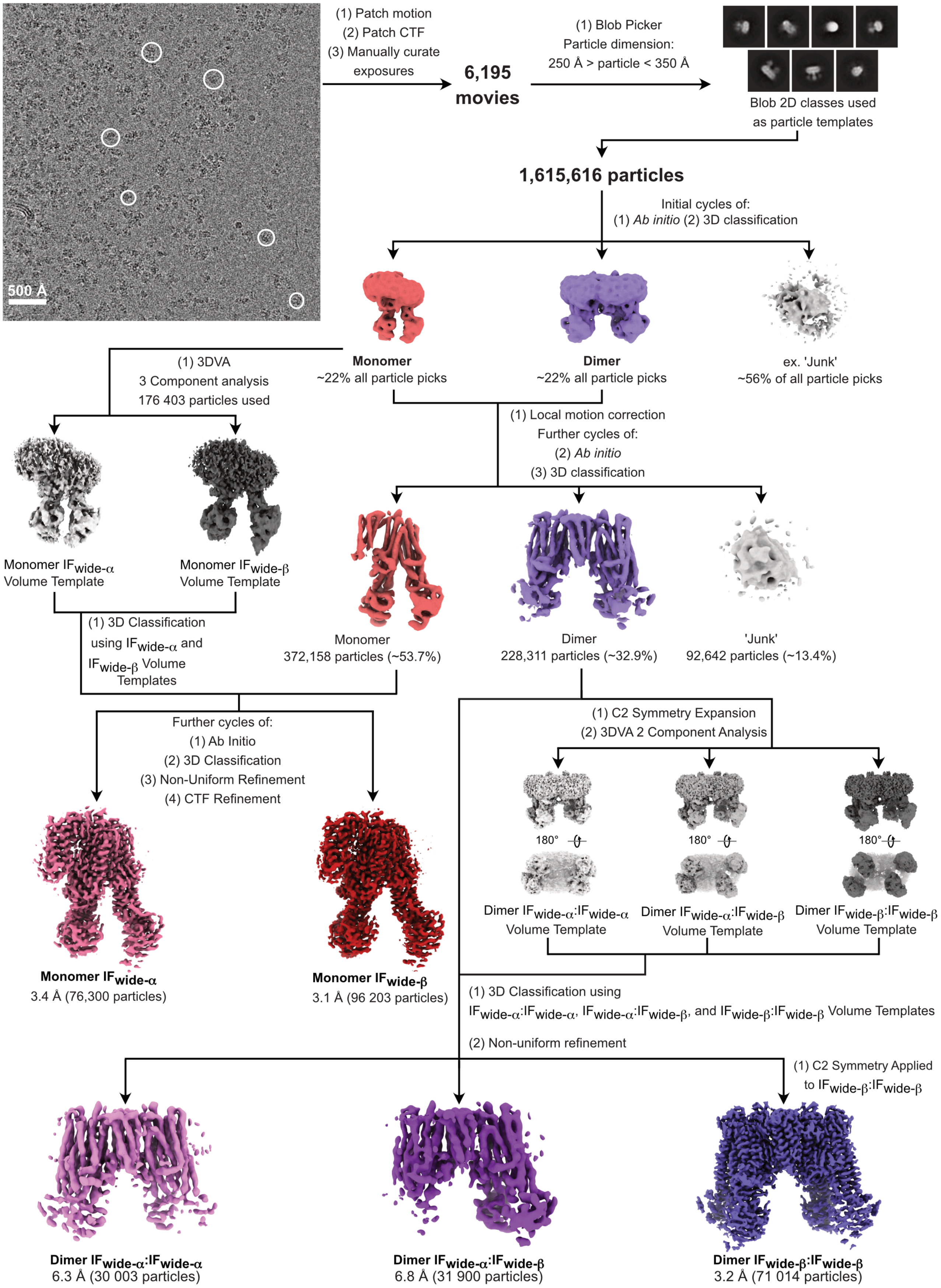


**Extended Data Fig. 2. Data collection and image processing workflow for determination of the cleaved Ycf1p monomer and dimer structures.** The flow-chart for cryo-EM data collection and processing, and structure refinement is shown, starting with a section of a representative micrograph (*top left*) and representative 2D classes (*top right*). Example Ycf1p particles in the micrograph are highlighted with a white circle (*top left*). Scale bar, 500Å. Also included in the flowchart are number of particles at each stage, as well as steps taken.

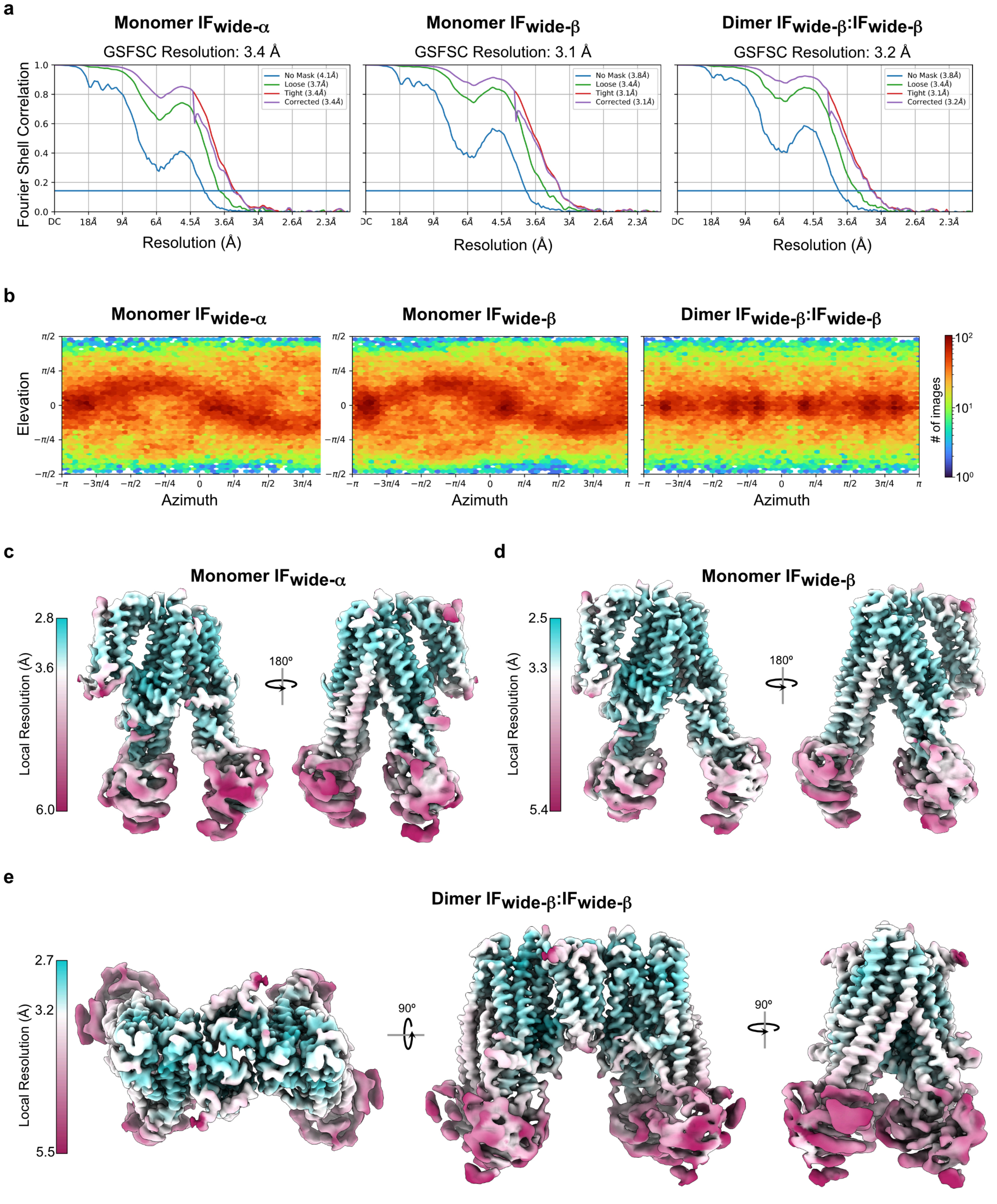


**Extended Data Fig. 3. Cryo-EM image analysis. a,** Fourier Shell Correlation (FSC) curves for the cryo-EM maps of the cleaved Ycf1p monomer in the IFwide-α (*left*) and IFwide-β (*centre*) conformations, and of the cleaved Ycf1p IFwide-β:IFwide-β dimer (*right*). **b,** Euler angle distribution for particle images contributing to each map. **c, d, e**, Local resolution estimated for the maps of the cleaved Ycf1p IFwide-α monomer (panel **c**), cleaved Ycf1p IFwide-β monomer (panel **d**), and the cleaved Ycf1p IFwide-β:IFwide-β dimer (panel **e**).

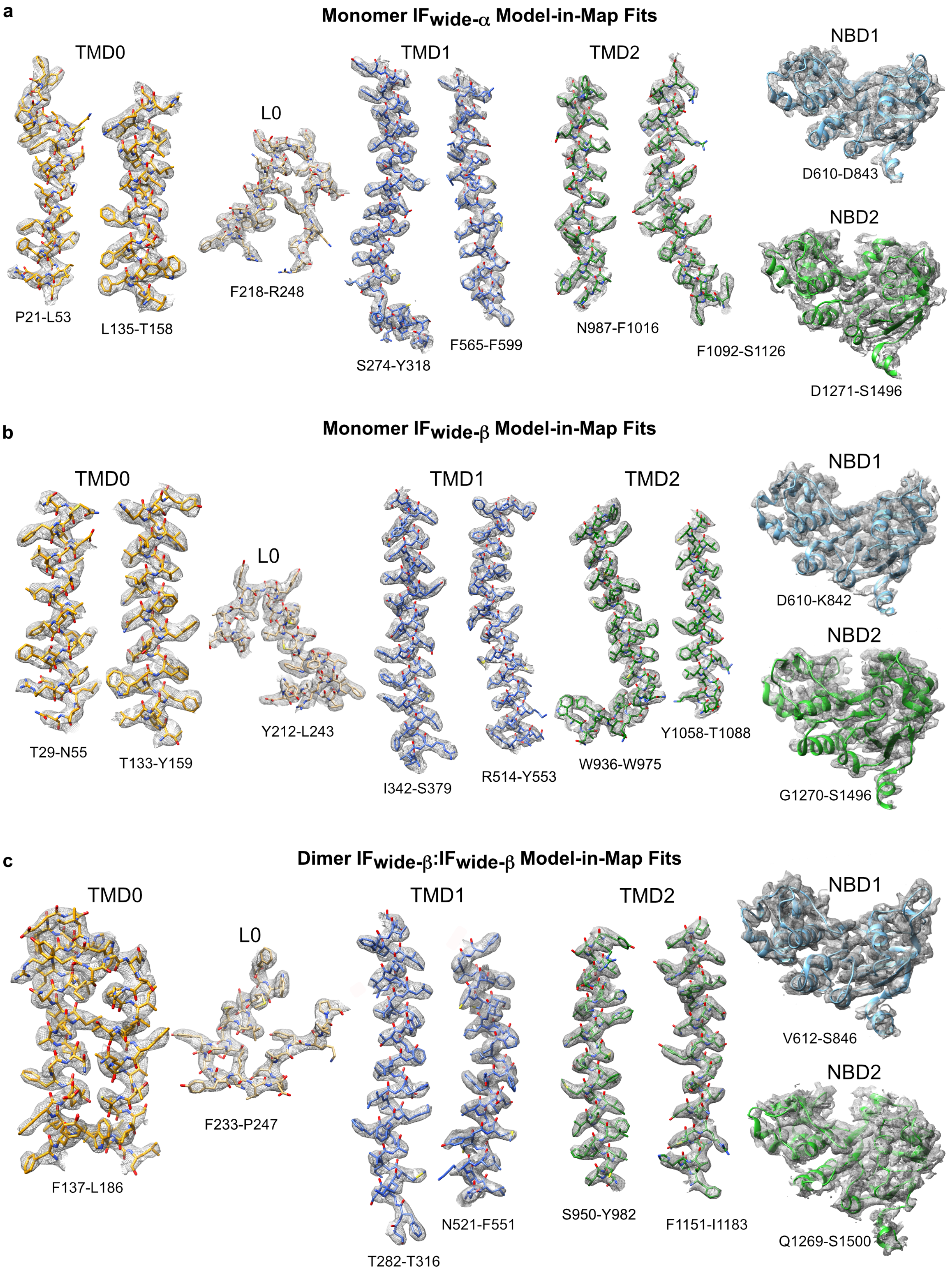


**Extended Data Fig. 4. Representative model-in-map fit for TMD0, the L0 linker, TMD1, TMD2, and the NBDs of the cleaved Ycf1p structures. a-c,** Examples of the atomic models of cleaved Ycf1p

IFwide-α monomer (panel **a**), cleaved Ycf1p IFwide-β monomer (panel **b**), and cleaved Ycf1p IFwide-β:IFwide-β dimer (panel **c**) fit into their respective experimental cryo-EM maps are shown for individual transmembrane helices in TMD0, TMD1, and TMD2, and various regions of the L0 linker. Schematic ribbon diagrams of NBD1 and NBD2 fit into the experimental cryo-EM maps, lacking any side chains, are shown on account of the fact that the NBDs are at lower resolution than the TMDs and L0 linker.

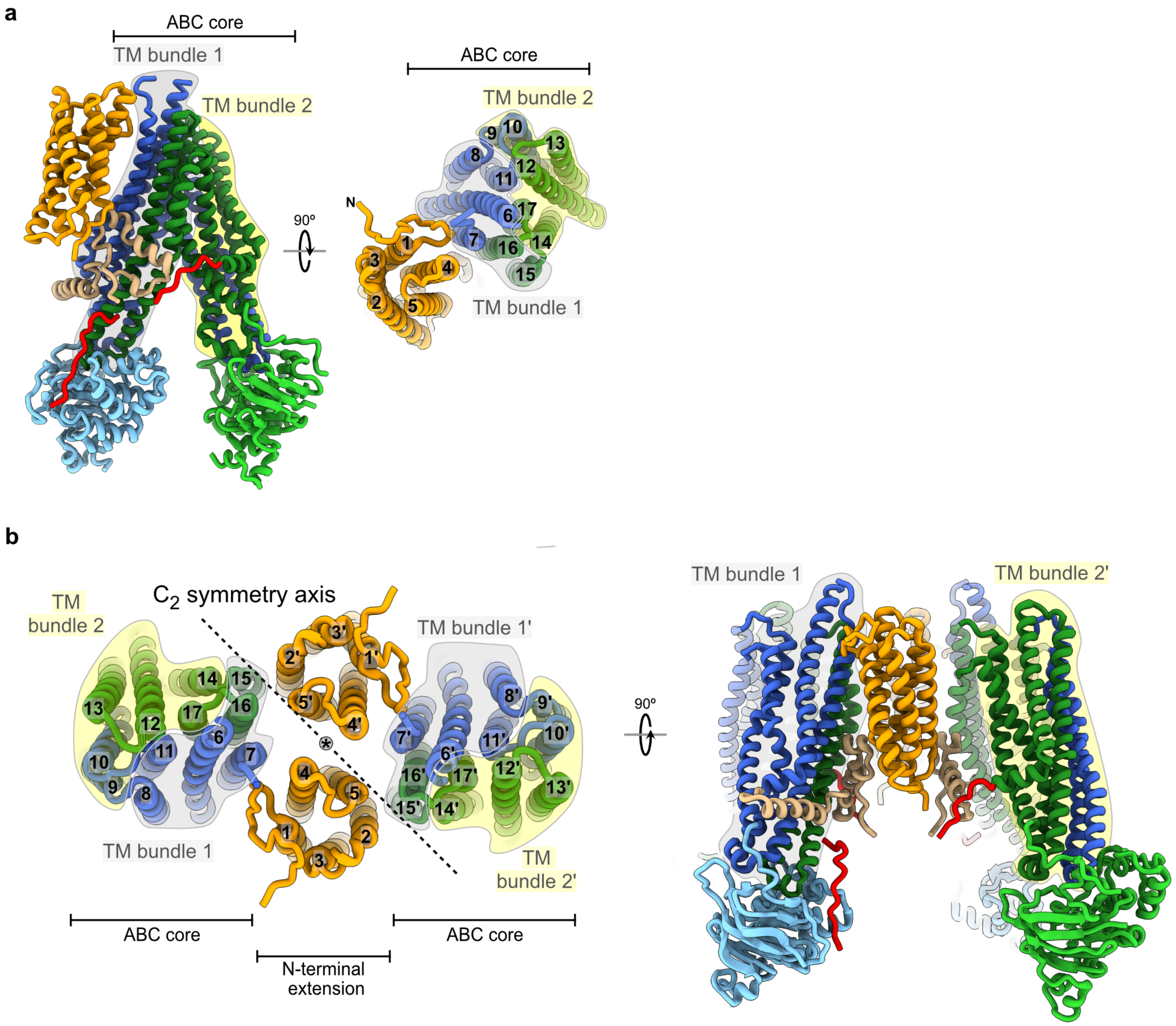


**Extended Data Fig. 5. Domain architecture of Ycf1p.** Schematic ribbon diagrams of the structures of the Ycf1p monomer (panel **a**) and the Ycf1p dimer (panel **b**). The top views, which are from the luminal side, highlight the composition of TM helix bundle 1 and TM helix bundle 2, which are encircled in grey and yellow, respectively. TM helices and TM helix bundles in one protomer of the Ycf1p dimer are labeled with a prime (“ ′ ”) in panel **b** to distinguish them from the opposite protomer.

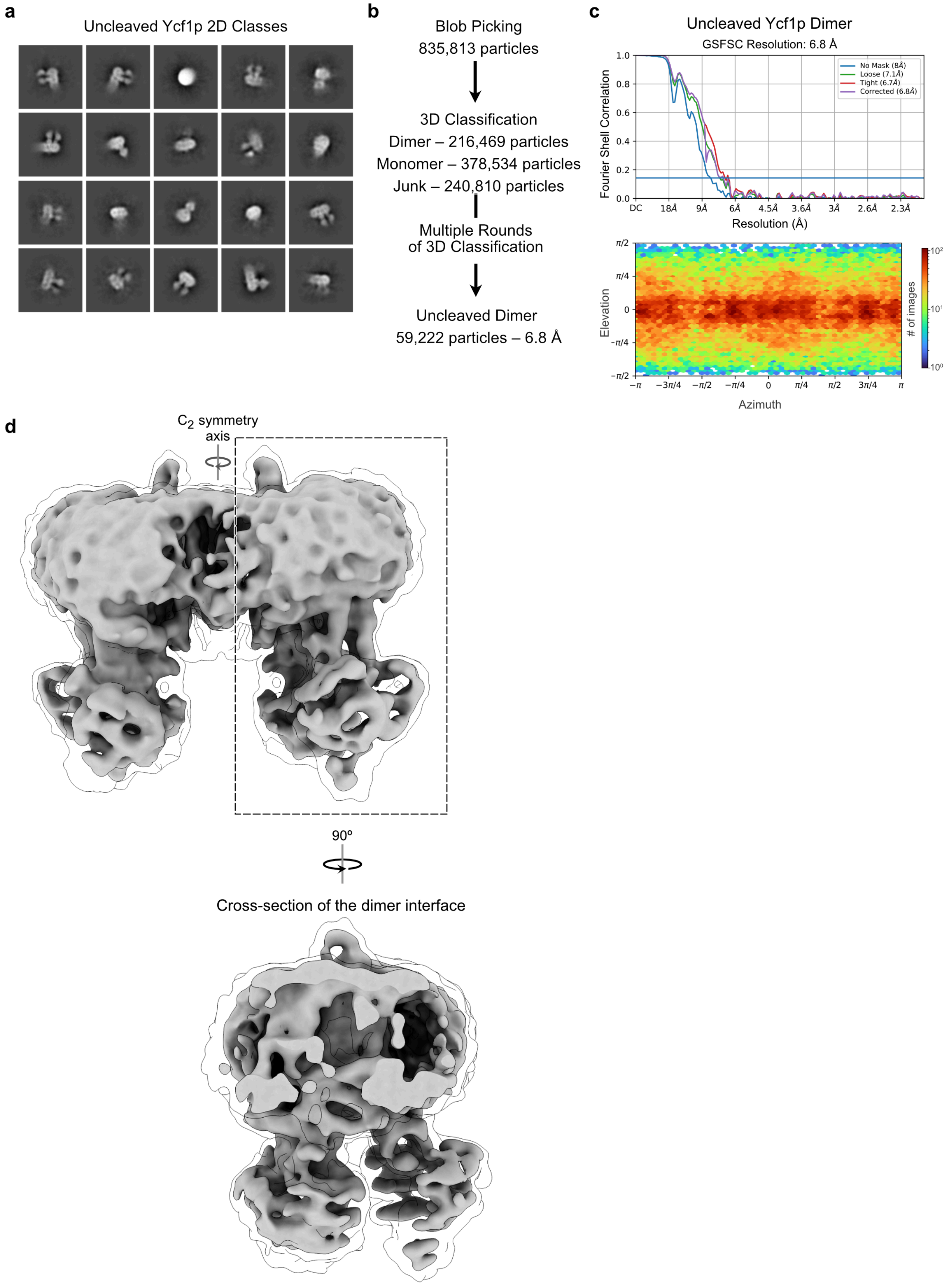


**Extended Data Fig. 6. Low-resolution map of an uncleaved Ycf1p dimer. a,** Representative 2D classes obtained from a previous cryo-EM data of uncleaved Ycf1p.[1] **b,** Flow-chart for analysis of uncleaved Ycf1p in which particles were picked based on the dimensions of the Ycf1p dimer. **c,** FSC curves for the uncleaved Ycf1p dimer (*top*) and Euler angle distribution for particle images contributing to the uncleaved Ycf1p dimer map (*bottom*). **d,** Cryo-EM density for the uncleaved Ycf1p dimer (6.8Å resolution), including a cross-sectional view of the dimer interface. The cryo-EM map lacks tubular density at the dimer interface that would correspond to TMD0 helices, and is further evidence that uncleaved Ycf1p does not adopt well-ordered and stable dimer.

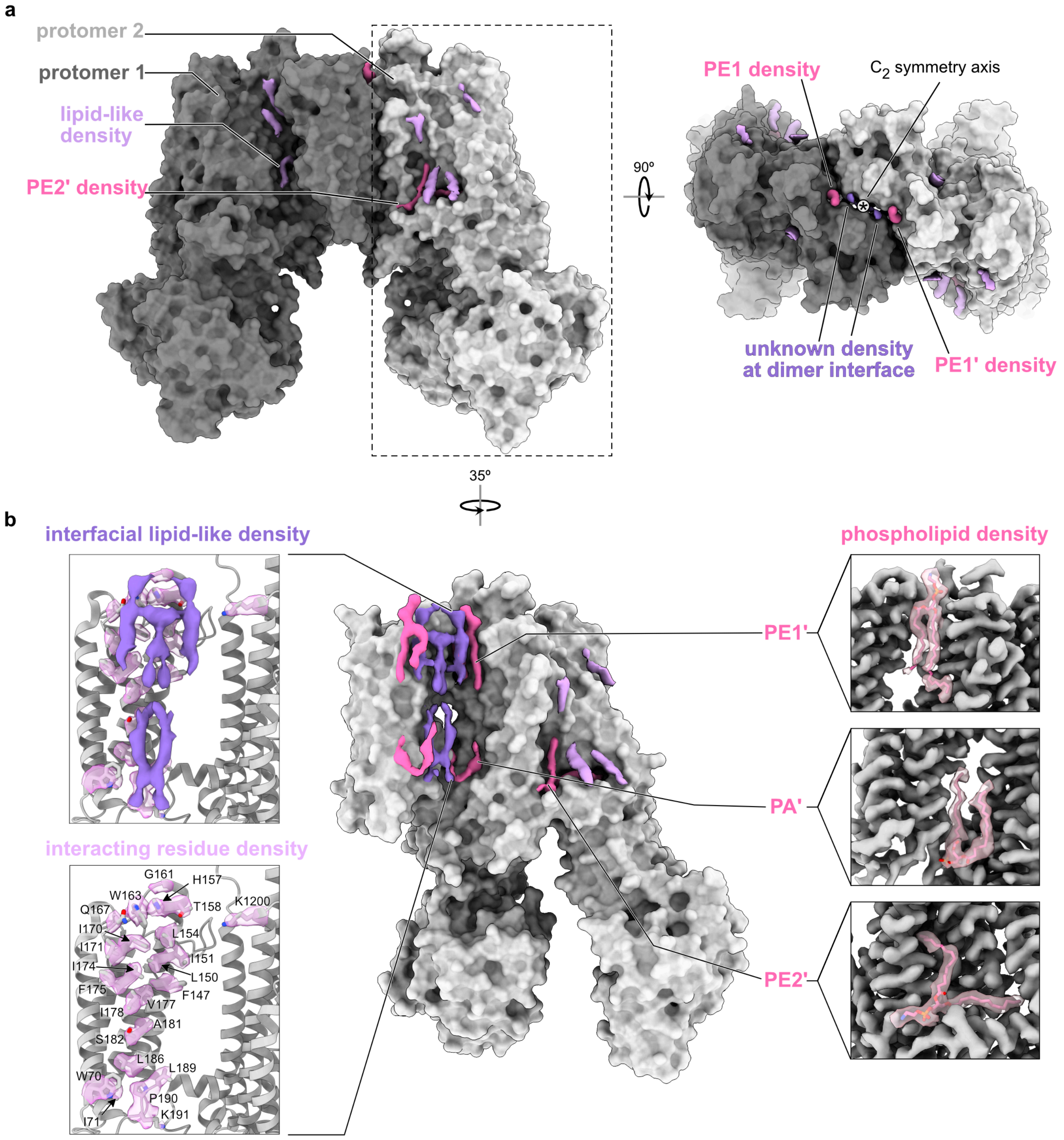


**Extended Data Fig. 7. Lipid densities in the transmembrane region of the Ycf1p dimer. a,** The cryo-EM map of the cleaved Ycf1p dimer is shown, coloured in dark grey for one protomer and light grey for the opposite protomer. The cryo-EM densities for bound lipids are shown in dark pink for the phosphatidylethanolamine (PE1, PE2) and the phosphatidic acid (PA) molecules that could be modeled, or in light or dark purple for bound lipids that could not be modeled. **b,** The cryo-EM map of one protomer and the associated lipid is shown with close-up views showing model-in-map fits of the phosphatidylethanolamine (PE1) bound at the luminal side between TMD0 and TM helix bundle 1 (*top right*), the interfacial phosphatidic acid bound between the two TMD0s on the cytosolic side of the dimer (*middle right*), and the phosphatidylethanolamine that contacts the R region and TMD2 in one protomer only (*bottom right*). Close-up views of the lipid density (dark purple) located in between the TMD0s in the Ycf1p dimer (*top left*). Cryo-EM density for interacting residues is shown (*bottom left*). The lipid density is removed from the lower panel to highlight all interacting residues, from one protomer, and for clarity. The identical residues from the opposite protomer also interact with these hydrophobic molecules.

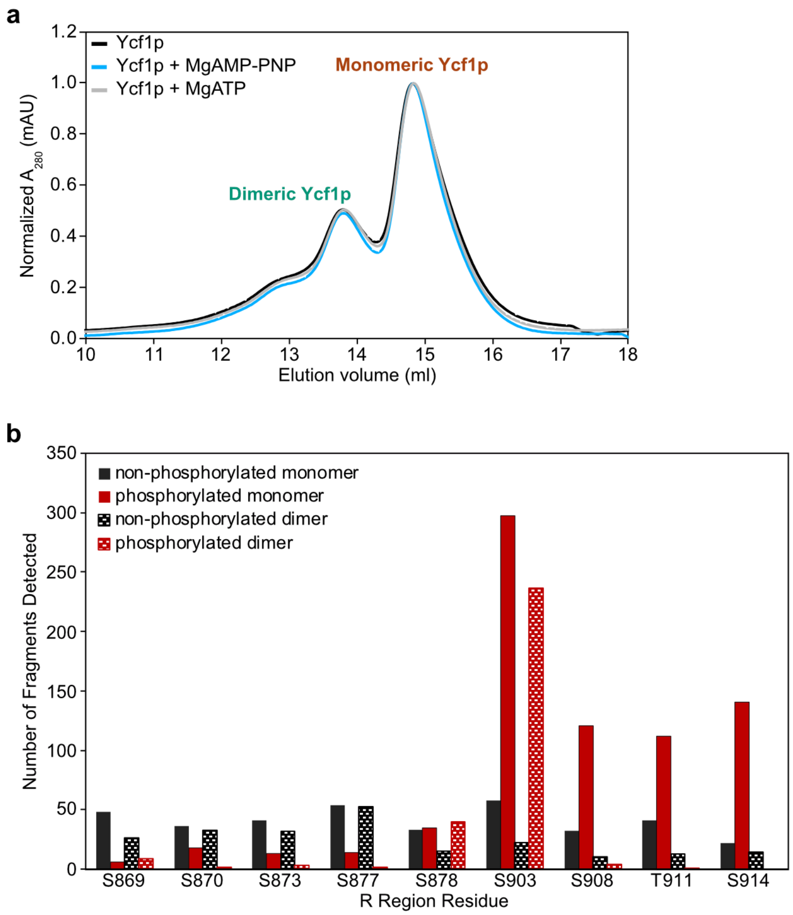


**Extended Data Fig. 8. Size exclusion chromatography and phosphorylation of cleaved Ycf1p. a,** The cleaved Ycf1p monomer and dimer populations are not altered by nucleotides, as shown by analytical gel filtration chromatography (Superose 6) of cleaved Ycf1p that was pre-incubated at 4 °C for 3-5h in the absence of nucleotides, in the presence of 2 mM $MgCl_2$ and 2 mM AMP-PNP, or in the presence of 2 mM $MgCl_2$ and 2 mM ATP. In each case, 130 µl of 0.81 mg/ml of protein was used. **b,** The monomeric and dimeric forms of cleaved Ycf1p display different levels of phosphorylation (right). Histogram displaying number of fragments from phosphorylated (red) and non-phosphorylated (black) peptides covering the indicated residue, as detected by LC-MS/MS. The number of fragments from monomeric Ycf1p are shown in solid bars, whereas the fragment count for dimeric Ycf1p are shown as hatched bars.
