## Supplementary Table 1 for "Structure of a dimeric ABC transporter"

| | Cleaved Ycf1p | | |
|---|---|---|---|
| | **Monomer** **$IF_{wide-\alpha}$** | **Monomer** **$IF_{wide-\beta}$** | **Dimer** **$IF_{wide-\beta}$:$IF_{wide-\beta}$** |
| **Data Collection** | | | |
| Electron microscope | Titan Krios G3 | | |
| Camera | Falcon4 | | |
| Voltage (kV) | 300 | | |
| Electron exposure (e-/Å$^2$) | 46.6 (1st data collection), 46.0 (2nd data collection) | | |
| Pixel size (Å) | 1.03 | | |
| Total exposure time (s) | 6.97 (1st data collection), 8.04 (2nd data collection) | | |
| Particle collection and density refinement software | cryoSPARC Live, cryoSPARC v2 | | |
| Combined movies (no.) | 6,195 | | |
| **Data Processing** | | | |
| Symmetry imposed | C1 | C1 | C2 |
| Initial particles (no.) | 1,615,616 | | |
| Final particles used in map (no.) | 76,300 | 96,203 | 71,014 |
| Global map resolution (Å) | 3.4 | 3.1 | 3.2 |
| FSC threshold | 0.14 | 0.14 | 0.14 |
| Map resolution range (Å) | 2.8-6.0 | 2.5-5.4 | 2.7-5.5 |
| **Model Building and Refinement** | | | |
| Initial model used | Monomer $IF_{wide\ \beta}$ | PDB 7MPE[1] | 2 x Monomer $IF_{wide\ \beta}$ |
| Ligands | PE: 1 | PE: 2 | PE: 4, PA:2 |
| Modelled residues | C10-P196, T205-D324, P337-L825, R926-P1263, G1270-C1506 | C10-H197, T205-D324, P337-L825, R929-P1263, G1270-C1506 | C10-P196, T205-H326, H336-L825, E927-P1262, Q1269-E1508 |
| Modelling and refinement software | Coot, Phenix, iSOLDE | | |
| R.M.S. bond length (Å) | 0.007 | 0.005 | 0.009 |
| R.M.S. bond angle (º) | 1.067 | 1.033 | 0.958 |
| Ramachandran outliers (%) | 0.00 | 0.00 | 0.00 |
| Ramachandran allowed (%) | 4.57 | 2.93 | 2.67 |
| Ramachandran favoured (%) | 95.43 | 97.07 | 97.33 |
| Rotamer outliers (%) | 0.00 | 0.16 | 0.00 |
| All-atom clashscore | 10.97 | 7.45 | 8.19 |
| MolProbity score | 1.88 | 1.57 | 1.57 |
